# Cryogenic electron tomography reveals novel structures in the apical complex of *Plasmodium falciparum*

**DOI:** 10.1101/2022.09.14.508031

**Authors:** Stella Y. Sun, Li-av Segev-Zarko, Grigore D. Pintilie, Chi Yong Kim, Sophia R. Staggers, Michael F. Schmid, Elizabeth S. Egan, Wah Chiu, John C. Boothroyd

**Author notes:** These authors contributed equally (Stella Y. Sun, Li-av Segev-Zarko). Correspondence (J.C.B), (W.C.).

## Abstract

Intracellular infectious agents, like the malaria parasite, *Plasmodium falciparum*, face the daunting challenge of how to invade a host cell. This problem may be even harder when the host cell in question is the enucleated red blood cell. Evolution has provided *P. falciparum* and related single-celled parasites within the phylum Apicomplexa with a collection of organelles at their apical end that mediate invasion. This apical complex includes at least two sets of secretory organelles, micronemes and rhoptries, and several structural features like apical rings and a putative pore through which proteins may be introduced into the host cell during invasion. In this paper, we perform cryogenic electron tomography (cryo-ET) on isolated merozoites to visualize the apical machinery. Through tomography reconstruction of cellular compartments, we see new details of known structures like the rhoptry tip interacting directly with a rosette resembling the recently described rhoptry-secretory-apparatus (RSA), or with an apical vesicle docked beneath the RSA. Subtomogram averaging reveals that the apical rings have a fixed number of repeating units, each of which is similar in overall size and shape to the units in the apical rings of tachyzoites of *Toxoplasma gondii*. Comparison of these polar rings in *Plasmodium* and *Toxoplasma* parasites also reveals them to have a structurally conserved assembly patterning. These results provide new insight into the essential features of this remarkable machinery used by apicomplexan parasites to invade their respective host cells.

## Introduction

*Plasmodium falciparum* is the causative agent of human malaria and, as such, represents a pathogen of global importance. Despite enormous efforts to control it, this apicomplexan parasite continues to infect > 220 million people each year, resulting in the deaths of ~627,000 individuals per year, mostly children under the age of five (WHO, 2021). One of the challenges in controlling this on-going pandemic is the rapidity with which drug-resistance emerges, making the discovery of new drugs and new drug targets an urgent task.

Like other Apicomplexa, *P. falciparum* is an obligate intracellular, single-celled parasite. Infections begin with introduction of sporozoites through the bite of an infected mosquito. These migrate to the liver where they infect hepatocytes, yielding several thousand infectious merozoites that are released into the bloodstream and go on to infect mature red blood cells (RBCs). After RBC invasion, the internalized merozoite undergoes a well-defined developmental cycle in which first the ring stage, then trophozoites, and then schizonts are produced over ~ 48 hours, culminating in the release of upwards of 30 daughter merozoites from each infected RBC (Cowman, Healer, Marapana, & Marsh, 2016). These in turn can go on to infect additional RBCs, producing waves of parasitemia. In this study, we focus on the merozoites, the small, invasive stage that transits between RBCs, for direct cryogenic-electron tomography (cryo-ET) imaging without sectioning or milling, which is usually required for thick cellular samples >1um (Rigort et al., 2012).

Invasion by merozoites is an active process driven mostly by the parasites themselves (Cowman, Tonkin, Tham, & Duraisingh, 2017; Paul, Egan, & Duraisingh, 2015). The first step is attachment through proteins present on the parasites’ surface binding to molecules on the RBC surface. This interaction can occur seemingly at almost any portion of the parasite’s and host’s respective surfaces. There soon follows, however, a reorientation of the parasite such that its apical end, containing key invasion machinery, is positioned in contact with the RBC. Next, the bulb-shaped secretory organelles known as rhoptries inject the contents of their apical necks into the host cell. These rhoptry neck proteins, or “RONs”, interact with host cytoskeletal components providing a key part of the machinery necessary for invasion. One RON protein, PfRON2, integrates into the RBC plasma membrane, forming a bridge between the RBC surface and its underlying cytoskeleton (Lamarque et al., 2011). PfRON2 binds a protein on the merozoite’s surface, PfAMA1, which is also an integral membrane protein, in this case originating from a different set of secretory organelles known as micronemes; these deposit their protein contents onto the merozoite’s surface through a mechanism whose details are not well understood but presumed to involve fusion of micronemes with the parasite’s plasma membrane. Once PfRON2 and PfAMA1 have bound one another, internalization can commence.

The rhoptries are secretory, bulb-shaped organelles that in *Plasmodium* are present as a pair, with their tapered ends oriented toward the merozoite’s apical end. The two rhoptries can also be found to be fused at their necks or go through complete fusion with each other during invasion (Bannister, Mitchell, Butcher, & Dennis, 1986; Hanssen et al., 2013). Their contents, which include many important effectors in addition to the RON proteins described above, are electron dense by transmission electron microscopy using conventional staining and fixation methods. Micronemes, which are also generally apical in placement, are much smaller than rhoptries and enriched at the apical region around the rhoptry. In addition to these two secretory organelles, the apical complex for which the phylum is named includes a set of anterior rings, the precise number depending on the species and the imaging being used. The most basal of these in *Toxoplasma* tachyzoites is the “apical polar ring” (APR) which is attached to microtubules and is therefore thought to function as a microtubule-organizing center (Koreny et al., 2021; NICHOLS & CHIAPPINO, 1987; Russell & Burns, 1984). The ring-like structures that have been described apical to the APR have been referred to in *P. falciparum* as apical rings (Hanssen et al., 2013; Koreny et al., 2021), in *P. berghei* as conoidal rings (Koreny et al., 2021), or in *T. gondii* as preconoidal rings (NICHOLS & CHIAPPINO, 1987). For simplicity, and given that in the *Plasmodium* merozoites there is no conoid cage and intraconoidal microtubules, we will refer to them here simply as “apical rings” (AR). The precise composition of these apical rings and their role in invasion or other biological functions of the merozoite are not yet known.

To help shed light on the interactions and possible functions of all these apical components, we have applied recently developed methods of cryo-ET to study merozoites embedded in vitreous ice without chemical fixation and staining. We report here the results of this examination, including details on the rhoptries and additional membrane bound organelles localized at the apical complex and new insight into the complexity of the apical ring structures.

## Results

### Cryo-ET of isolated merozoites reveals new details of the apical complex

*P. falciparum* (strain 3D7) was grown in human blood using established methods, and synchronized using successive rounds of sorbitol and magnetic purification. Cultures at the early schizont stage were incubated with the protease inhibitor E-64 for 5-6 hours, and then free merozoites were harvested using syringe filtration as previously described (Boyle et al., 2010). After washing the harvested merozoites, they were concentrated and loaded onto a Lacey carbon grid, followed by one-side blotting and plunge-freezing in liquid ethane. This freezing process avoids artifacts caused by chemical fixation or staining used for conventional TEM sample preparation. The frozen grids were transferred to a 200KV Talos Arctica electron microscope and imaged with Volta Phase Plate optics and sample-stage tilting. Each tilt image followed a step of 2- or 3-degree increments in order to compute a 3-D volume of the sample. A representative tomogram of the whole merozoite cell is presented in video 1. Virtual sections from this tomogram and other representative tomograms are presented in Fig. 1.

**Fig. 1.**
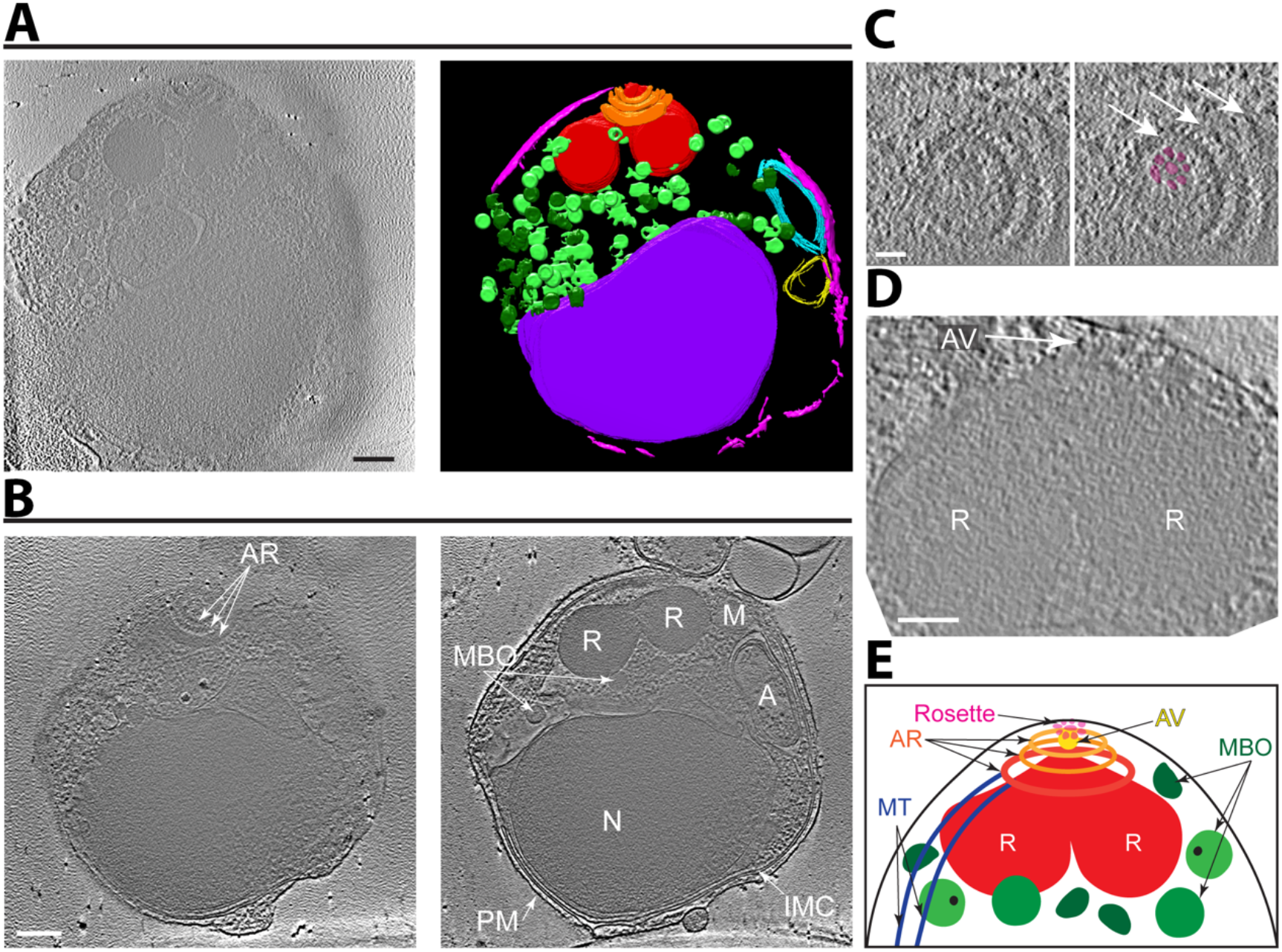
Cryo-ET reveals the organization of subcellular organelles in *Plasmodium falciparum* merozoites. (**A**) Tomographic slice of a representative merozoite (left) and its 3-D annotation showing the apical rings (orange), rhoptries (red), membrane-bound organelles (light/dark green), inner-membrane complex (pink), apicoplast (cyan), mitochondrion (yellow), and nucleus (purple). Scale bar, 200 nm. Movie with the complete tomogram and annotation is in Video 1. (**B**) Two tomographic slices of a merozoite (different from that shown in (A)) better showing the apical rings (AR), rhoptries (R), membrane-bound organelles (MBO), inner membrane complex (IMC), apicoplast (A), mitochondrion (M), nucleus (N), and plasma membrane (PM). Scale bar, 200 nm. (**C**) Zoomed-in view of a tomographic slice of a merozoite (different from (A) and (B)) showing the apical rings (arrows) and the rosette (highlighted in magenta). Scale bar, 50 nm. (**D**) Zoomed-in view of a tomographic slice of a merozoite (different from (A), (B), and (C)) showing the apical vesicle (AV; arrow) at the tip of two rhoptries (R). Scale bar, 100 nm. (**E**) Cartoon of the *Plasmodium* merozoite apical complex showing the arrangement of key subcellular structures.

Given the limited penetrating capability of an electron beam, cryo-ET is generally restricted to cellular samples < 0.5 microns. We were therefore surprised that our tomogram captured not only the tapering apical end, which was our primary objective, but also the entire length and width of many parasites that normally are roughly spherical and about 1.5 microns in diameter. We attribute this to the parasites being disrupted during sample preparation and cannot exclude the possibility that there was consequent loss of some of their contents. Nevertheless, we readily identified all the known cytoskeletal structures and organelles of the merozoite, including their apical rings with associated microtubules, a pair of rhoptries, membrane-bound organelles that presumably are a mix of micronemes and dense granules, the inner membrane complex (IMC), the nucleus, the apicoplast with its multiple delimiting membranes, and mitochondria (Fig. 1). Our primary interest is in the components of the apical complex, which we and others have recently analyzed in detail in cryo-ET studies with the related apicomplexan parasite, *T. gondii* (Mageswaran et al., 2021; Sun et al., 2022; Wang et al., 2021); we therefore focused our attention on this area. Of particular note was the presence of a distinct rosette of particles at the extreme apical tip of the merozoites (Fig. 1C). This and other features are described in more detail below.

### Apparent densities associate with the membrane-bound organelles

In cells that appeared minimally disrupted, we identified, in addition to the rhoptries, three subclasses of membrane-bound organelles (MBO), distinguished by their shape and small size (Fig. 2). While the contents of two subclasses appeared relatively homogeneous, the third subclass of MBO had a very electron-dense cluster inside (Fig. 2A,B). Which, if any, of these corresponds to the micronemes or dense granules cannot be discerned from the data presented here but they seem likely to have very distinct cargoes and, likely, functions, particularly the third subclass.

**Fig. 2.**
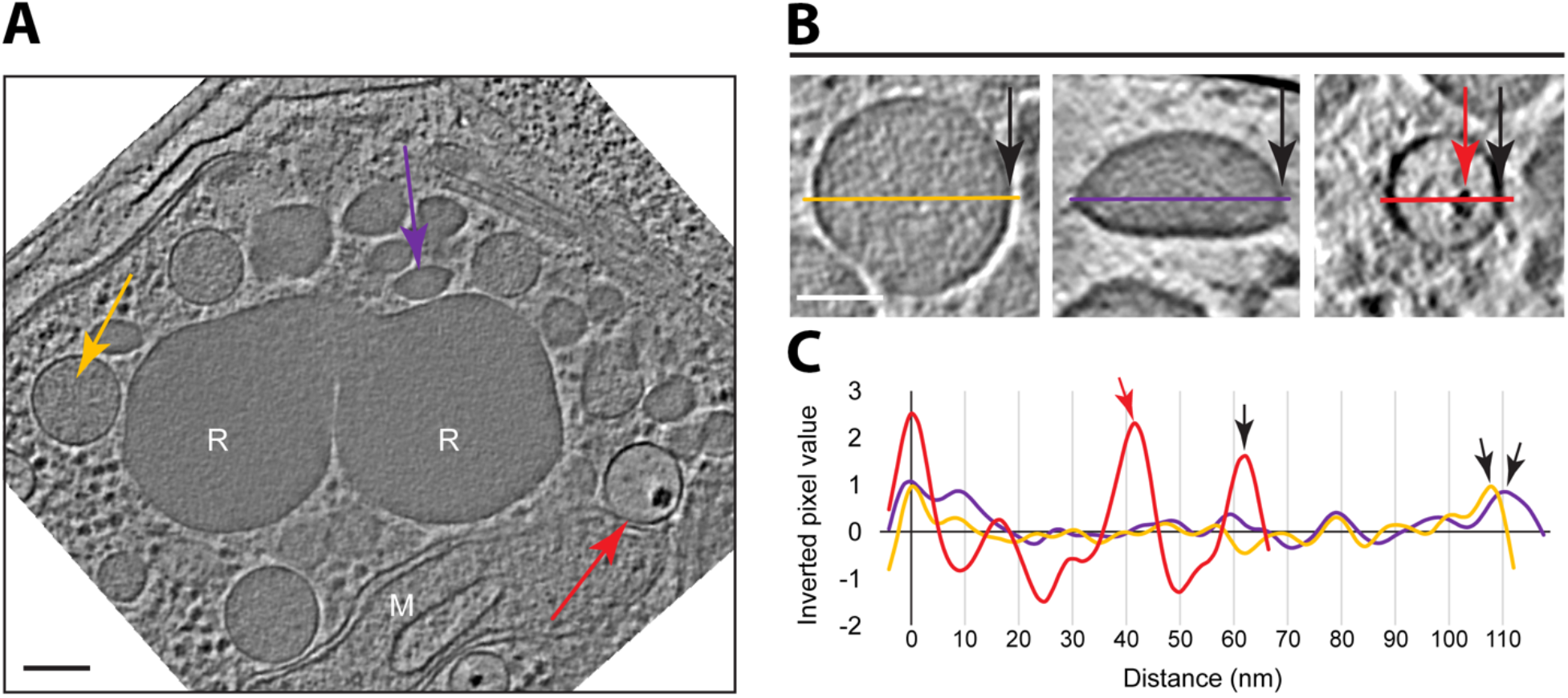
Three distinct subclasses of membrane-bound organelles are present. **(A)** A tomographic slice of a *Plasmodium* merozoite showing the rhoptries (R), mitochondria (M), and three distinct subclasses of membrane-bound organelles, MBO1, MBO2, and MBO3, marked with yellow, purple and red arrows, respectively. Scale bar, 100 nm. (**B**) Zoomed in views of a representative of each subclass, including a line colored as in (A) and with a black arrow marking the right edge in all cases. The red arrow indicates the electron-dense region characteristic of MBO3s. (**C**) Inverted pixel values of the lines shown in (B). Arrows also as in (B). Scale bar, 50 nm.

Close inspection of the rhoptry interior mostly revealed a granulated texture with no apparent subsections or ordered densities. Along the inner side of the rhoptry membrane, however, we identified an additional density resembling a lining (Fig. S1A). On the outer surface of this organelle, we observed scattered, discrete globular densities attached to only the neck area of the rhoptries, while the rest of the exterior surface of the rhoptry bulb appeared smoother (Fig. S1B-E). The identity and significance of these densities are not known.

In addition to the single-membrane-bound organelles described above, double- and quadruple--membrane-bound organelles were also observed (Fig. 1 and S2); these have all the very characteristic features, and are therefore presumed to be, the mitochondrion and apicoplast, respectively. When both structures were detected in the same tomographic slice, they were always observed next to one another, as previously reported (Hopkins et al., 1999; van Dooren et al., 2005) (Fig. S2). The shortest distance between the outer membrane of the mitochondrion and the apicoplast was similar to the distance between the inner and outer membranes of the mitochondria or the two outermost membranes of the apicoplast in the same area (Fig. S2B). A close inspection revealed densities in the interface between the mitochondria and the apicoplast in the areas of intimate association, suggesting a discrete structure connects them (Fig S2C,D).

### A rosette is apparent on the merozoite surface with an apical vesicle beneath it

We observed a distinct density pattern at the apical end of these merozoites, consisting of a rosette embedded within the parasite’s plasma membrane and concentric with the apical rings (Fig. 3, video 2). Each rosette has an 8-fold rotational symmetry with an overall diameter of ~70 nm and also has an axial ring-like density (Fig. 3A). A similar rosette-like structure has been described at the apical end of *Toxoplasma* tachyzoites and *Cryptosporidium* sporozoites, and has been termed the rhoptry secretory apparatus (RSA) because of its presumed role in secretion by these organelles (Mageswaran et al., 2021). In 14 out of 35 tomograms, we identified an apical vesicle immediately beneath the rosette (Fig 3A, Video 2). The shape of the AVs was often less spherical than the AVs observed in *Toxoplasma* with an average diameter of 46.6 ± 4.2 nm (N=14). In instances where the vesicle was missing, the tip of the rhoptries appeared to directly interact with the rosette (Fig. S3A, B, C and Video 3). This interaction was evident even in cases where the plasma membrane integrity was compromised, and the rhoptries were separated from their position close to the apical rings. This condition resulted in an elongated structure that appeared to be membrane-limited and that extended all the way up into and through the space surrounded by the apical rings, eventually reaching the merozoite surface (Fig. S3D and Video 3). Although the origin of this membrane cannot be definitively discerned from these images, the data are consistent with it being an extended invagination of the parasite’s plasma membrane, perhaps a result of the rhoptries being firmly attached to the membrane (through the RSA) and pulled away during sample preparation.

**Fig. 3.**
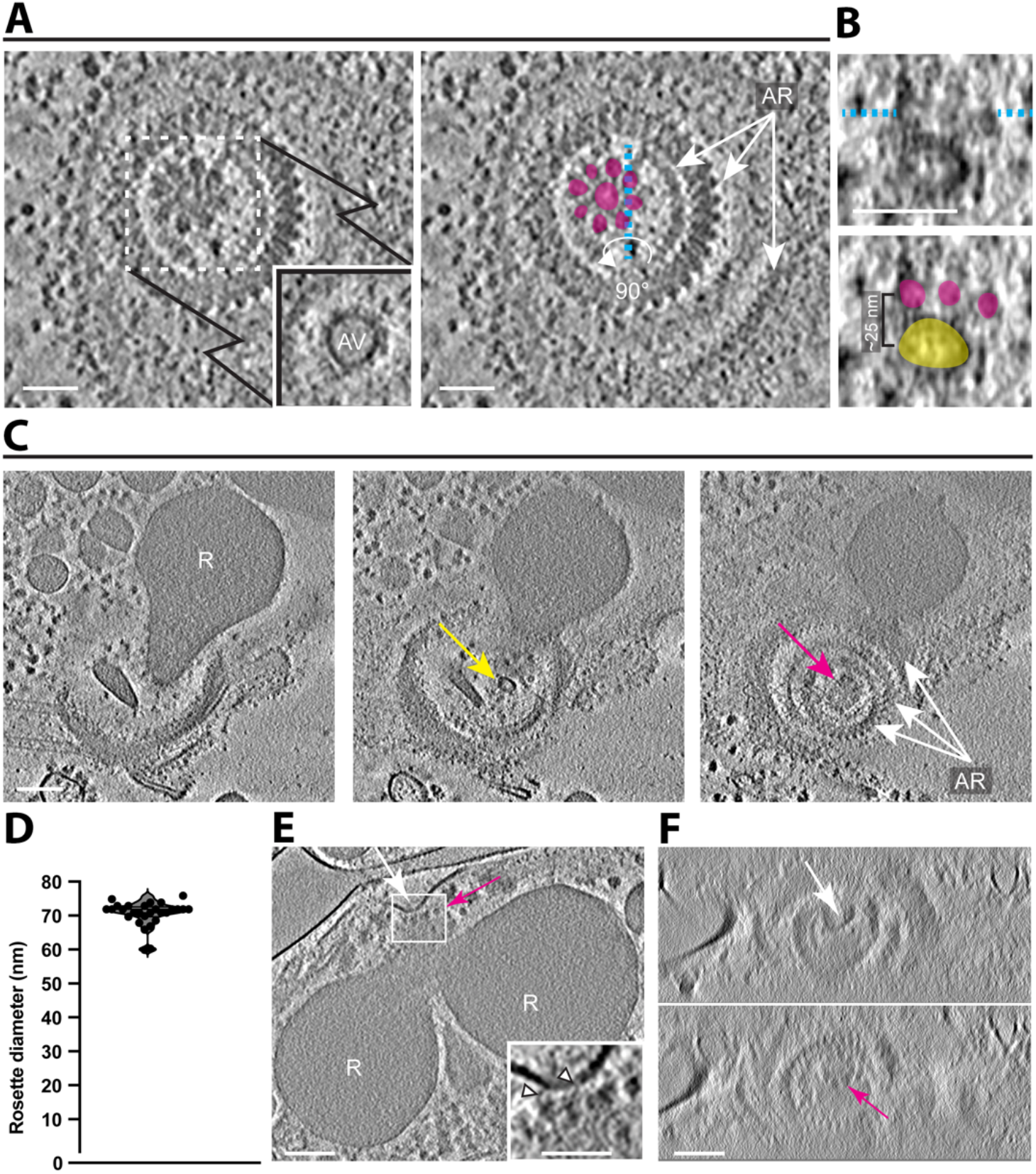
Rosettes are associated with either an apical vesicle or a rhoptry tip. (**A**) A tomographic slice showing a top view of the three apical rings (AR) surrounding a rosette. The solid-line square shows an apical vesicle (AV) in a z-slice just below the area indicated by the dashed-line square. Right panel highlights the densities of the rosette in magenta. Scale bar, 50 nm. (**B**) A zoomed-in side view of the rosette and the AV from (A) showing the orthogonal sectioning plane labeled in cyan dashed line. Bottom panel highlights the rosette in magenta and the AV in yellow. (**C**) Three tomographic slices from a single tomogram (different from (A)) showing a single rhoptry (R) with an AV at its tip (yellow arrow) and above it a rosette (magenta arrow) surrounded by the three ARs. Scale bar, 100 nm. (**D**) Rosette diameters (between outer edges) as measured from the top view of the rosette (mean ± SD =70.7±3.7 nm, N=28). (**E**) A tomographic slice showing a dimple in the parasite membrane (white arrow) in the area of association with the rosette (seen from a side view, magenta arrow) and the two rhoptries (R). Scale bar, 100 nm. A zoomed in view of the rectangle shows densities between the parasite membrane and the rosette. Scale bar, 50 nm. (**F**) An orthogonal view of the tomogram in (E) showing the top-down view of the dimple in the parasite membrane (top panel, white arrow) and the rosette in the z-slices just below (bottom panel, magenta arrow). Images are a stack of 7 consecutive slices. Scale bar, 100 nm.

### The apical rings present a prominent 34-fold symmetry

Three generally distinct, concentric rings were apparent at the apical end of the parasites, with diameters of ~180, ~240 and ~370 nm shown in slice view, respectively (Fig. 4A, D and F). To gain structural details on the smallest, most apical ring as well as the next closest ring (“ring 1” and “ring 2,” respectively), we performed subtomogram averaging on extracted particles, each representing 5 repeating units. The results showed substantial mass in both rings with a bridge density connecting the two (Fig. 4B). Note that distinct beads like densities were visible on the inner-most edge of ring 3 (Fig. 4A), but no ordered mass was resolved in the subtomogram averaged map (Fig. 4B). The resulted averaged density was mapped back to the cellular tomograms to assemble the complete rings 1 and 2 and to determine the precise number of repeating units from top view (Fig. 4C). The results showed a highly reproducible, 34-fold symmetry based on the horizontal slice view and 3D volume of the cryoET density map from 27 out of 39 tomograms, indicating this is the number of units comprising both rings (Fig. 4C, S4 and Video 4). Although fewer repeating units were observed in some tomograms, the difference is likely a result of the partial or off-angle view that did not allow a complete image of the entire ring. In some cases, the rings appeared bent into somewhat distorted circles, suggesting some flexibility in the connection between each ring unit (Fig. S6). Ring 1 can be further segmented into an inner ring (pink) with a diameter of ~125.6 nm and an outer ring density (red) with a diameter of ~186.8 nm (Fig. S3B and S4C). The spacing between the neighboring units of outer ring 1 is ~17.3 nm (Fig. S4B). The segmentation of ring 2 revealed a circular density of the inner ring (blue) associated with 34-fold rotational density (green) with a diameter of ~232 nm and a ~21.7 nm neighbor spacing (Fig. 4D, Fig. S3C and S4B,C). Most importantly, rings 1 and 2 can be seen to be clearly connected via a “bridge” (Fig. S3A; yellow) which also presents as 34 repeating units, connecting the bottom, outer edge of ring 1 and the inner, upper edge of ring 2. Thus, these two rings are clearly connected.

**Fig. 4.**
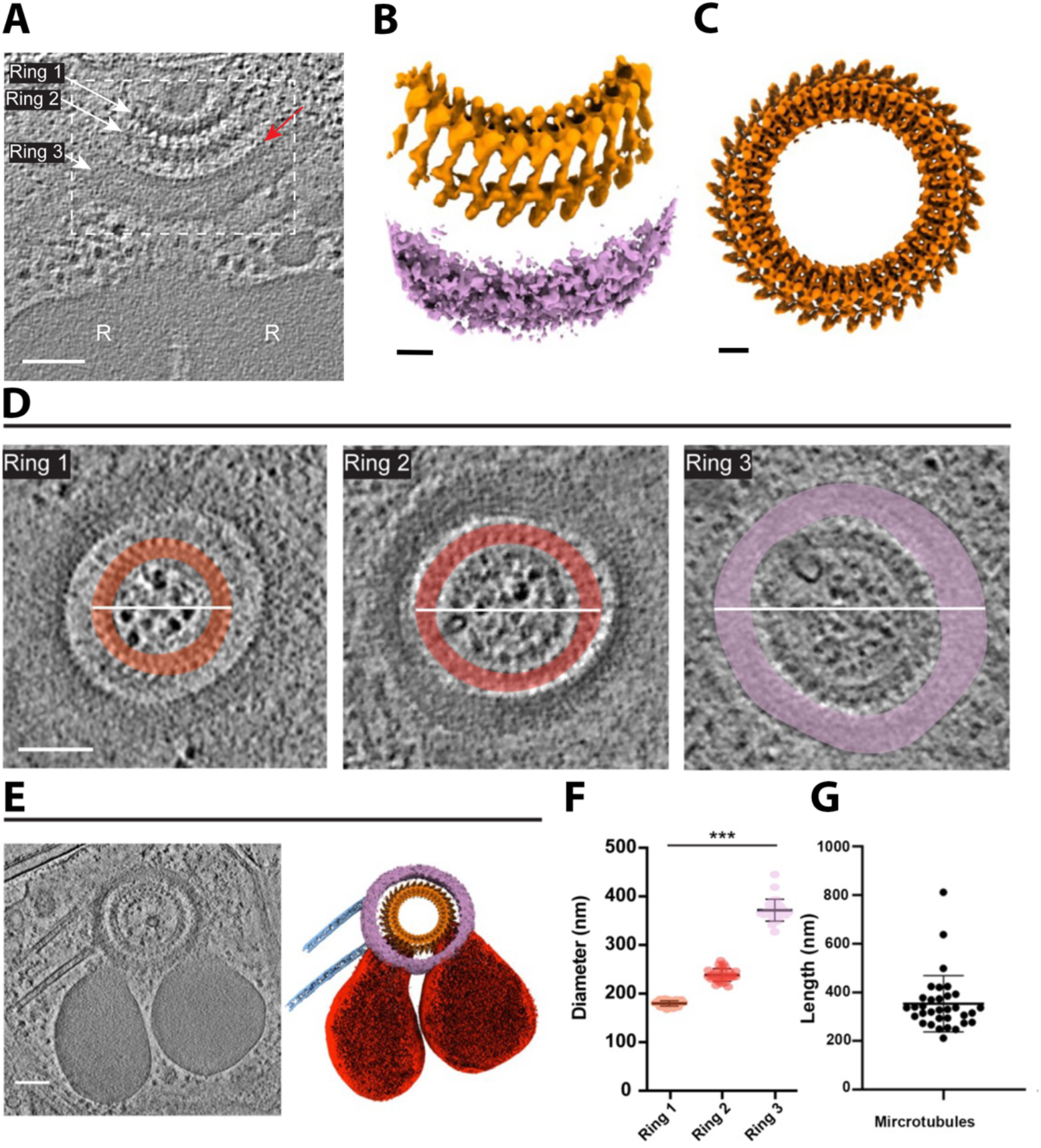
The apical rings appear as three distinct entities. (**A**) A tomographic slice showing the rhoptries (R) and three apical rings marked Ring 1, 2, and 3, respectively. Red arrow points to “beads” like densities on the inner-most edge of ring 3. The dashed square shows the approximate region of the apical rings represented in (B). Scale bar, 500 nm. (**B**) A subtomogram averaged map of the dashed box area shown in (A). Scale bar, 20 nm. (**C**) The average map of rings 1 and 2 in (B) is mapped back to a cellular tomogram to assemble their entirety and shown from a top view. Scale bar, 25 nm. (**D**) Three tomographic slices from a tomogram (different from (A)) showing a top view of the three apical rings, highlighted in different colors (orange, red, and violet for Rings 1, 2, and 3, respectively). The white lines represent the diameter that was measured for (F). Scale bar, 100 nm. (**E**) Left panel is a tomographic slice showing the rhoptries, three apical rings, the apical vesicle, and two subpellicular microtubules (SPMTs). Right panel is the cryo-ET average densities of the three apical rings (gold and purple) and the annotation of SPMTs (blue) and rhoptries (red). Scale bar, 100 nm. (**F**) The mean maximum diameter for each apical ring. Ring 1 mean ± SD =180±6 nm, N=30; Ring 2 mean ± SD =238±13 nm, N=30; Ring 3 mean ± SD =372±23 nm, N=29. (**G**) The mean length of associated microtubules is 353.8±115.5 nm, N=33.

Using ring 2 and the bridge portion as anchor points, focus refinement was applied to the third ring, resulting in a cryo-ET density map that appeared relatively amorphous, had no obvious repeating features, and was about 50 nm thick (Fig. 4B). One or two subpellicular microtubules were anchored to the edge of ring 3 in 21 tomograms (N=39) with a length ranging between 212.1 nm and 812.2 nm for any one microtubule (mean ± SD =353.8±115.5 nm, N=33) (Fig. 4G).

Focus refinement was further applied to rings 1 and 2 to reveal more features of structural densities for each repeating unit (Fig. 5). The focused refinement map of rings 1 and 2 was fitted into the whole ring structure (Fig. 5A). Based on the vertical repeating pattern, a small, distinct unit spanning both rings can be further segmented into the entire structure (Fig. 5B,C). Based on the density volume and using prior calibration to the tubulin subunits of microtubules in *Toxoplasma*, the total mass of the repeating unit of this ring complex can be estimated to be between 13.6 MDa and 16.3 MDa.

**Fig. 5.**
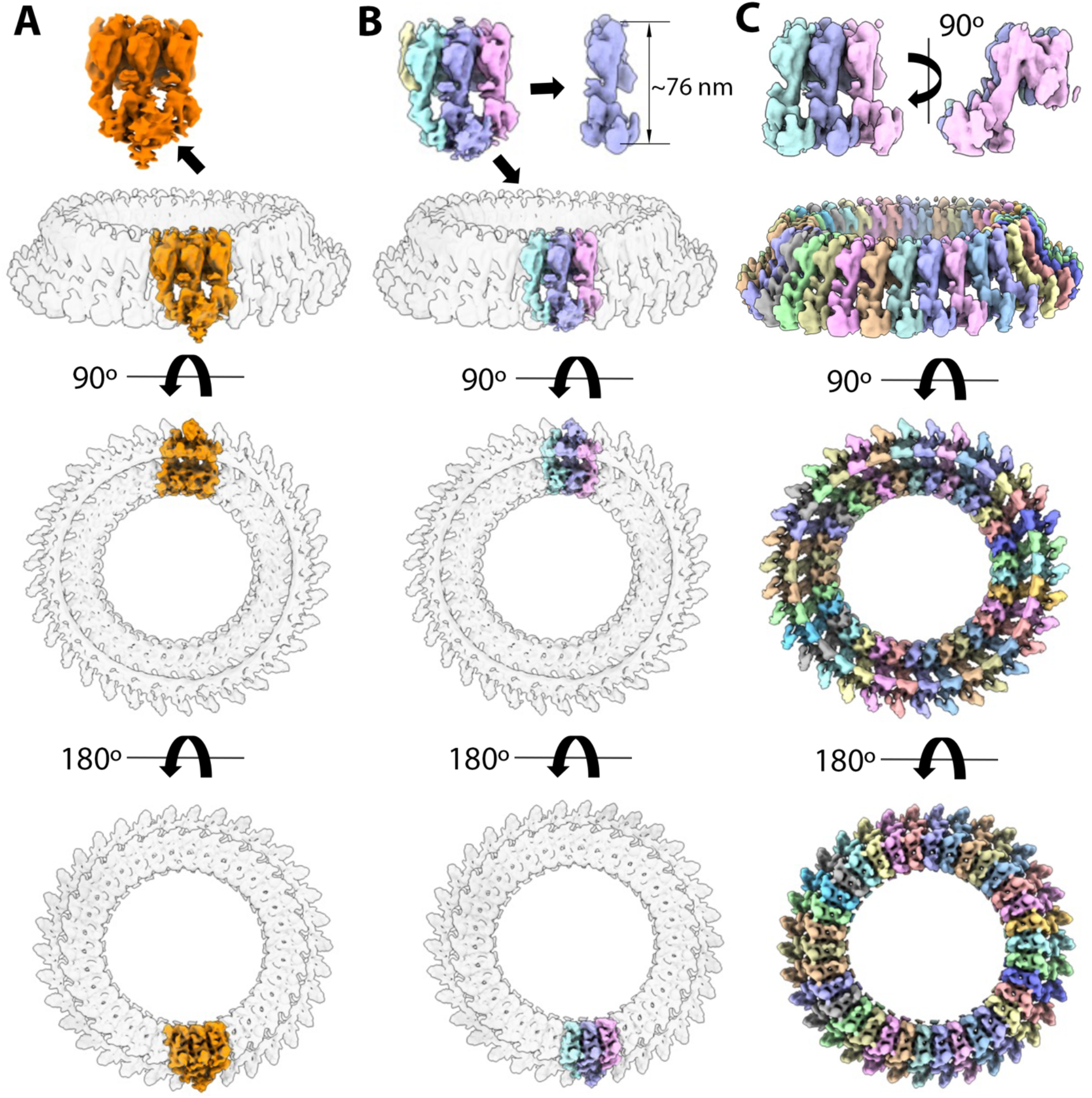
Rings 1 and 2 of *Plasmodium falciparum* merozoites contain 34 repeating units. (**A**) Averaged map of rings 1 and 2 by focused refinement was fitted and aligned with the whole circular ring. (**B**) Based on the repeating pattern of densities, the repeating unit of the apical ring segment can be further segmented into 3 repeating units shown in cyan, blue and pink color. (**C**) Using the complete repeating unit (blue) as a template, 34 repeating units of the rings were segmented and shown in different colors.

The subtomogram averaged map was then used to compare the overall density of rings 1 and 2 of *Plasmodium* merozoites with that of the repeating units averaged from the preconoidal rings of *Toxoplasma* tachyzoites (Fig. 6 and Video 5). The overall dimension of the *Toxoplasma* preconoidal rings is ~220 nm in maximum outer diameter shown in slice view and their overall arrangement was similar in being two concentric and somewhat discrete rings with a bridge linking the two. When the bridge densities were aligned and fitted between *Toxoplasma* and *Plasmodium*, they are oriented in different directions. However, they can be locally aligned (Fig. 6B). In addition, the averaged maps reveal a ~76 nm distance between the Plasmodium top and bottom apical rings while the distance between the *Toxoplasma* rings is ~65 nm. When aligned, ring 2 of the *Toxoplasma* preconoidal rings extends ~21 nm lower than the *Plasmodium* ring 2 (Fig. 6B). The space between two successive densities bridging the two rings is about ~15.5 nm and ~17.7 nm for *Plasmodium* and *Toxoplasma*, respectively (Fig. 6B). In addition, the *Toxoplasma* repeating units were considerably more in number; although we were not able to determine a precise count, we observed ~42-45 units in five tomograms where they could be estimated. Importantly, the overall size and shape of the repeating units segmented from rings 1 and 2 of *Toxoplasma* and *Plasmodium* are similar (Fig. 6C and D).

**Fig. 6.**
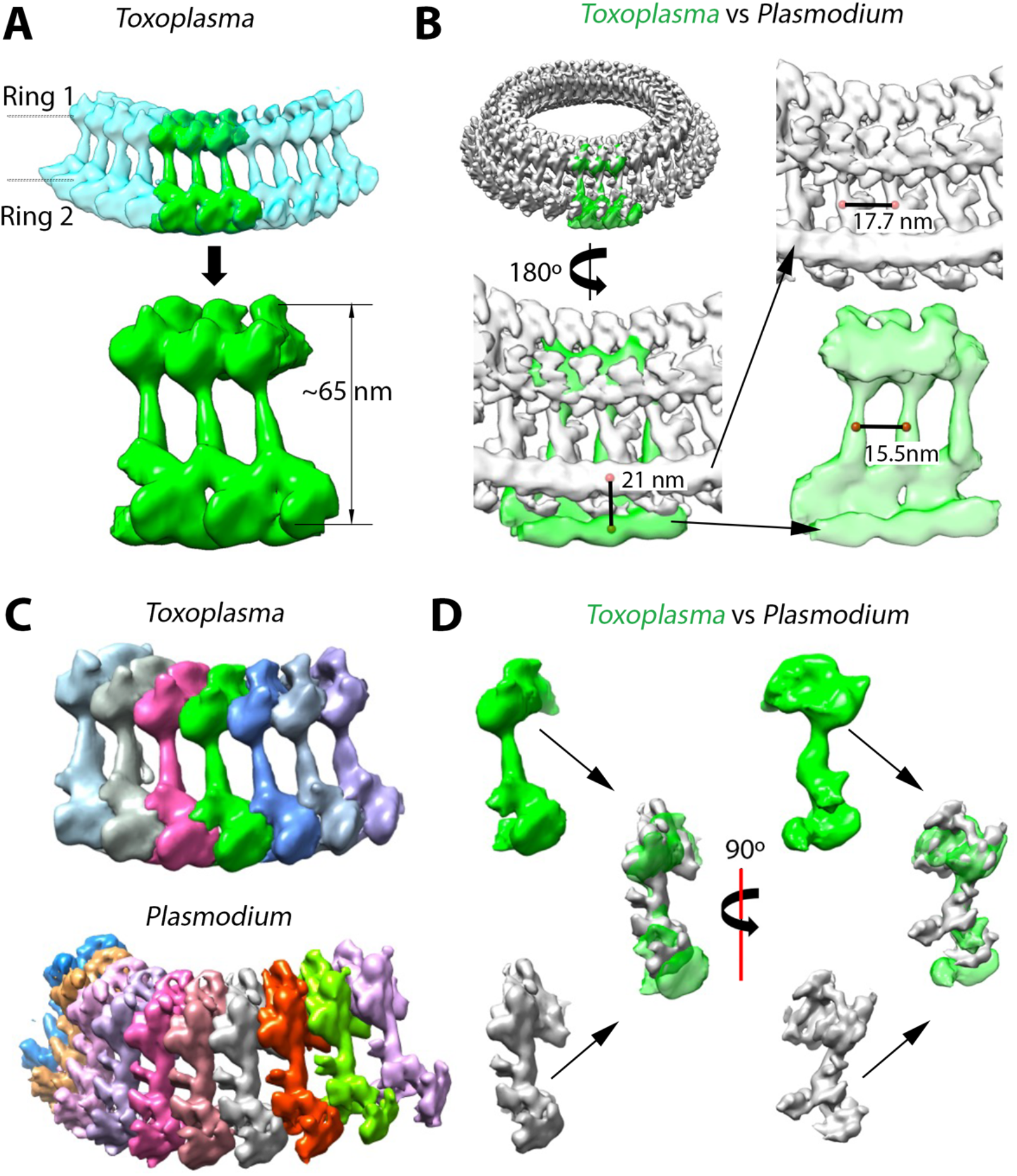
*Plasmodium falciparum* rings 1 and 2 show a similar repeating unit and pattern to those seen for the pre-conoidal rings of *Toxoplasma gondii*. (**A**) The averaged map of the repeating ring units of *Toxoplasma* tachyzoites (data from previously published work; (Sun et al., 2022)) can be mapped back to a cellular tomogram to show the assembled ring shown in light sea blue. A segment containing 3 repeating units (green) is also shown expanded. (**B**) The *Toxoplasma* segment from (A) was fitted and aligned into the averaged map of apical rings 1 and 2 of *Plasmodium* (gray). A portion of the resulting alignment is shown after 180º rotation, i.e., as viewed from the inside of the rings. Note that ring 2 extends lower in *Plasmodium* than in *Toxoplasma* by ~21 nm, and the spacing between neighboring units is smaller in *Toxoplasma* (~15.5 nm) compared to *Plasmodium* (~17.7 nm). (**C**) A single repeating unit in the average map of apical polar rings was further segmented from *Toxoplasma* and *Plasmodium*. (**D**) The comparison of a single unit from *Plasmodium* and *Toxoplasma* rings shows structural similarities from the front (on the left) and side views (90° counterclockwise viewing from top, on the right).

## Discussion

We report here images of the apical ends of vitrified *P. falciparum* merozoites, revealing intriguing structures not previously described. One of the structures observed is a rosette which extends from the tip of a rhoptry neck or apical vesicle adjacent to just below the parasite membrane. Given their size, pattern and position, we presume these are analogous, and likely homologous, to the RSA recently described in the related *Toxoplasma* and *Cryptosporidium* (Mageswaran et al., 2021). As shown by those authors, this structure appears to have a common evolutionary origin with the discharge apparatus of the very distantly related ciliates like *Paramecium*. As for the Apicomplexa, these are members of the superphylum Alveolata and it has been proposed that orthologues of the well-characterized “Nd” proteins of ciliates comprise one or more portions of the rosette in *Toxoplasma. Plasmodium*, too, has orthologues of these same Nd6 and Nd9 proteins (Amos et al., 2022; Aquilini et al., 2021; Mageswaran et al., 2021) and so it seems highly likely that these serve the same function here.

In both *Plasmodium* merozoites and *Toxoplasma* tachyzoites, freeze-fracture microscopy has revealed the presence of a small, external pit or dimple at the center of the parasite’s apical tip (Aikawa, Miller, Rabbege, & Epstein, 1981; Dubremetz, 2007). This structure is at the center of the rosette mentioned above and has been speculated to function as part of the channel connecting the parasites’ rhoptries with the host cell cytosol. In this study we were not able to determine if the rosette might be the source of this surface structure but the center tip of the rosette lies at about the position of the dimple and so it seems likely that they are related in function, if not physically connected.

In many tomograms, we also detected a small, apical vesicle sandwiched between the rhoptry tip and the rosette, supporting the recent reports (Martinez et al., 2022; Segev-Zarko et al., 2022). The location and lower density of this vesicle (relative to the rhoptries) are both reminiscent of the apical vesicles seen in *Toxoplasma* tachyzoites (Mageswaran et al., 2021; Paredes-Santos, De Souza, & Attias, 2012; Segev-Zarko et al., 2022). In the latter parasite, the most apical of these vesicles undergoes an apparent fusion with the tip of the rhoptry necks when the parasites are exposed to a calcium ionophore, a stimulus that causes *Toxoplasma*’*s* conoid to be protruded in preparation for invasion (Segev-Zarko et al., 2022). Many have speculated that this apical vesicle is part of the machinery that enables rhoptries to secrete their contents into the host cell through pores at the start of tachyzoite invasion (Mageswaran et al., 2021), some even referring to such as “porosomes” (Monteiro, De Melo, Attias, & De Souza, 2001; Paredes-Santos et al., 2012). It is similarly tempting to speculate that the rosette of *Plasmodium* might also serve as a conduit for moving their rhoptry contents into a red blood cell.

Although the precise composition of the apical rings is not known, some components have been described (Dos Santos Pacheco, Tosetti, Koreny, Waller, & Soldati-Favre, 2020; Hu et al., 2006; Koreny et al., 2021); their precise location within the rings, however, has yet to be determined. Our subtomogram averaging shows a clear linkage between ring 1 and ring 2 with a bridge density connecting the two. This is similar to the overall organization of the preconoidal ring in *Toxoplasma* tachyzoites but the *Plasmodium* ring had substantially fewer subunits comprising it (34 vs. ~42-45 for *Plasmodium* and *Toxoplasma*, respectively). In addition, the curvature changes of the *Plasmodium* ring didn’t break the connection of the repeating ring units, suggesting a resilience to the rings that may play a role in the invasion process when encountering physical barriers within the host cells. It seems likely that the respective compositions of the ring complexes in *Plasmodium* and *Toxoplasma* will be similar but much work remains to be done on the exact proteins involved and their relative locations within each ring.

## Materials and Methods

### *Plasmodium falciparum* culture and preparation of free merozoites

*P. falciparum* strain 3D7 was routinely cultured in de-identified human erythrocytes from the Stanford Blood Center at 2% hematocrit at 37°C, 5% CO_2_ and 1% O_2_. The culture medium (termed complete RPMI) consisted of RPMI-1640 (Sigma) supplemented with 25 mM HEPES, 50 mg/L hypoxanthine (Sigma), 2.42 mM sodium bicarbonate, and 4.31 mg/ml Albumax II (Gibco).

Free merozoites were isolated as previously described by Boyle et al., with some modifications (Boyle et al., 2010). *P. falciparum* was synchronized to the schizont stage using sequential rounds of 5% sorbitol (Sigma) and magnet purification using LS columns (Miltenyi Biotec) and a MACS magnet. On the day of harvest, 120 ml of *P. falciparum* culture at 10-20% parasitemia were subjected to MACS magnet purification to isolate schizonts away from uninfected erythrocytes, yielding ~5×10^8^ schizonts. The schizonts were resuspended in 40 ml fresh complete RPMI with 10µM E-64 (Sigma), a cysteine protease inhibitor that prevents merozoite release from schizonts by inhibiting rupture, but does not adversely affect merozoites (Boyle et al., 2010; Glushakova, Mazar, Hohmann-marriott, Hama, & Zimmerberg, 2009). After a 5 hour incubation at 37°C, the cells were pelleted at 1500 rpm for 5 minutes and supernatant removed completely. The schizont pellet was resuspended in 6ml of PBS supplemented with 2% FBS, 5 mM MgCl_2_, and 50 µg/ml DNase I, and transferred to a syringe connected to a 1.2 µm filter. To release the merozoites, pressure was applied to the syringe plunger evenly until all liquid was expelled. The flow-through was applied to an LS column on a MACS magnet to remove hemozoin, and then centrifuged at 4000 rpm for 10 minutes, followed by two washes in PBS. The merozoites were quantified using a hemocytometer, and resuspended at 5×10^7^/ml or 5×10^6^/ml.

### *Toxoplasma gondii* culture and preparation of free tachyzoites

*Toxoplasma gondii* RH*∆hxgprt* strain was maintained by growth in confluent primary human foreskin fibroblasts (HFFs) in Dulbecco’s modified Eagle’s medium (DMEM; Invitrogen, Carlsbad, CA) with 10% fetal bovine serum (FBS; HyClone, Logan, UT), 2 mM glutamine, 100 U/ml penicillin, and 100 μg/ml streptomycin (cDMEM) at 37°C in 5% CO2. The HFFs are fully deidentified and therefore do not constitute human subjects research.

*Toxoplasma* tachyzoites were released from heavily infected monolayers of HFFs by mechanical disruption of the monolayers using disposable scrapers and passage through a syringe attached to a 25-gauge needle. The parasites were added to fresh monolayers of HFFs, and 18-20 hours post-infection, were washed two times with cold Hank’s balanced salt solution (HBSS) without calcium, magnesium and phenol red (Corning, Corning, NY), supplemented with 1 mM MgCL_2_, 1 mM CaCl_2_, 10 mM NaHCO_3_ and 20 mM HEPES, pH 7. The HFF monolayers were scraped and passed through a 27-gauge syringe and tachyzoites were released into fresh cold HBSS. Tachyzoites were pelleted and resuspended in fresh cold HBSS and calcium-ionophore (A23187, Sigma) at a final concentration of 1 μM was added to the sample at room temperature.

### Cryogenic electron tomography

Free merozoites or tachyzoites suspension mixed with 10 nm gold fiducials (EMS) were loaded to the lacey carbon grids, blotted from the back side, and plunged into liquid ethane cooled down to liquid nitrogen temperature using Leica GP. The parasites were imaged using a Talos Arctica electron microscope (Thermo Fisher) equipped with a field emission gun operated at 200kV, a Volta Phase Plate (Danev, Buijsse, Khoshouei, Plitzko, & Baumeister, 2014), an energy filter (Gatan) operated at zero-loss and a K2 Summit direct electron detector (Gatan). Upon phase plate alignment and conditioning, the low-magnification tilt series of whole parasites were recorded at a magnification of 24,000x at pixel size 5.623 Å, and high-magnification tilt series of the apical region were recorded at a magnification of 39,000x at pixel size 3.543 Å, using Tomo4 software with bidirectional acquisition schemes, each from −60° to 60° with 2° or 3° increment. Target defocus was set to −1.0 µm. The K2 camera was operated in dose fractionation mode recording with 0.4 or 0.5 per e/Å^2^ frame. To increase the contrast, an energy filter was used with a slit width of 20eV, and a new spot on the phase plate was selected every 2 or 3 tilt series. The phase shift spans a range from 0.2-0.8π. The total dose was limited to 100-120 e/Å^2^ for merozoite samples, and 70-90 e/Å^2^ for tachyzoite samples.

### Tomography Reconstruction and Analysis

The movie frames were motion-corrected using motionCor2 (Zheng et al., 2017), and the resulting micrographs are compiled into tilt series. For *Plasmodium* merozoites, a total of 18 tomograms of whole merozoites at a magnification of 24,000x, and 39 tomograms at the apical region of merozoites at a magnification of 39,000x were collected. Tilt series alignment, tomography reconstruction, and CTF estimation are performed automatically using the tomography pipeline in EMAN2 (Chen et al., 2019). For features inspection and presentation, tilt series alignment and reconstruction was performed using IMOD (Kremer, Mastronarde, & McIntosh, 1996; Mastronarde, 1997). Subcellular features were manually segmented using Fiji and viewed in Chimera (Pettersen et al., 2004; Schindelin et al., 2012). The repeating units of apical rings were selected and centered manually in the middle of each repeating unit using EMAN2 (Chen et al., 2019). Subtomograms are then generated with an 80% overlap between neighboring ring units. De novo initial models are directly generated from the particles. For the *in situ* apical ring structure of merozoites, 1,156 particles from 39 tomograms were used in the subtomogram average. Starting from the successful subtomogram refinement of the apical ring unit, focused refinement was applied to the bottom part of the ring unit by extracting the particle with a large box size to include the MTOC ring. For the subtomogram averaging of the apical ring of *Toxoplasma* tachyzoites, 335 particles were generated from 59 tomograms collected with a magnification of 39,000x. The average map of the ring unit can be mapped back into tomograms for assembly of the circular whole rings in *Plasmodium* and *Toxoplasma*. Subsequent visualization and animation of the tomograms and maps were using Chimera or ChimeraX (Pettersen et al., 2004).

### Distance measurements

The distance between objects was measured using a line profile plot of the inverted pixel value. Distance between the apicoplast and mitochondria membranes was evaluated along a line crossing at the minimal distance between the organelles. The radii of the *Plasmodium* rings were measured by the outermost density for each of the 3 rings based on the slice views.

### Map Segmentation

All maps were segmented using the Segger plugin v2.9.1 in UCSF Chimera (Pintilie, Zhang, Goddard, Chiu, & Gossard, 2010). First, the map of the plasmodium ring was segmented into ring 1, bridge, and ring 2. Rings 1 and 2 also contain inner and outer sub-rings (as shown in Figure S4). The map of the *Plasmodium* apical rings (Figure 4) was segmented without using any smoothing and grouping, and the regions were grouped interactively to generate two regions representing rings 1 and 2 (orange) and ring 3 (purple). The densities for rings 1 and 2 were then segmented again, using one “smoothing and grouping” step of size 1, after which segments corresponding to repeating units were interactively grouped; each repeating unit consisted of ~20 segments. For the map of the *Toxoplasma* rings (Figure 6), a region with 8 repeating units was first cropped from the full map. The cropped map was segmented without using any smoothing and grouping, and again segments corresponding to repeating units were identified and grouped interactively; each repeating unit consisted of ~8 segments. For comparison purposes (Figure 6), the map of the cropped *Toxoplasma* rings was interactively aligned to the map of the full *Plasmodium* rings 1 and 2 in UCSF Chimera, and the alignment was refined using the Fit in Map dialog. Two possible alignments were obtained (depending on the starting alignment), one in which the main vertical bridge stalks are parallel, and the other in which the ring 2s from each map were parallel (Figure 6). The Volume Tracer in Chimera was used to place markers and measure distances between the stalk portions in adjacent repeating units, and also the difference in position of the base of ring 2 in *Toxoplasma* vs. *Plasmodium*. The molecular weights of repeating units in each organism (Figure S5) were estimated from the volume of each repeating unit measured using the “Measure Volume and Area” dialog in Chimera, assuming an even density throughout the units of 1.35e-24 g/A3 (Quillin & Matthews, 2000), and the approximate conversion units from kg to kDa of 1.660e-24. The radii of the *Plasmodium* rings were measured by interactively placing markers on the averaged map using the Volume Tracer and Distances dialogs in Chimera (Pettersen et al., 2004).

## Data availability

Representative tomograms of *Plasmodium falciparum* and subvolume averages of maps are deposited to EMDB. The accession code for the 3D reconstruction of *Plasmodium* free merozoite is EMD-28142. The accession codes for the reconstructions showing the apical complex of *Plasmodium* are EMD-28138 (side view) and EMD-28141 (top view). The accession codes for the *Plasmodium* averaged maps are EMD-28123 for the whole assembled ring1 and ring2, EMD-28124 for the ring1, ring2 and ring3 segment, and EMD-28125 for the repeating unit of ring1 and ring2 is. The accession codes for the *Toxoplasma* average map are EMD-28126 for ring1 and ring2 segment, and EMD-28127 for the repeating unit of ring1 and ring2.

## Acknowledgements

We thank the CZI Biohub Intercampus Team Award and Stanford ChEM-H for supporting this research. Other support includes NIH grants: R01GM079429, DP2HL137186, Vaadia-BARD Postdoctoral Fellowship Award No. FI-582-2018, the Stanford Maternal and Child Health Research Institute, and Stanford School of Medicine Dean’s Postdoctoral Fellowship.

