## Supplementary material for "Cryogenic electron tomography reveals novel structures in the apical complex of *Plasmodium falciparum*"

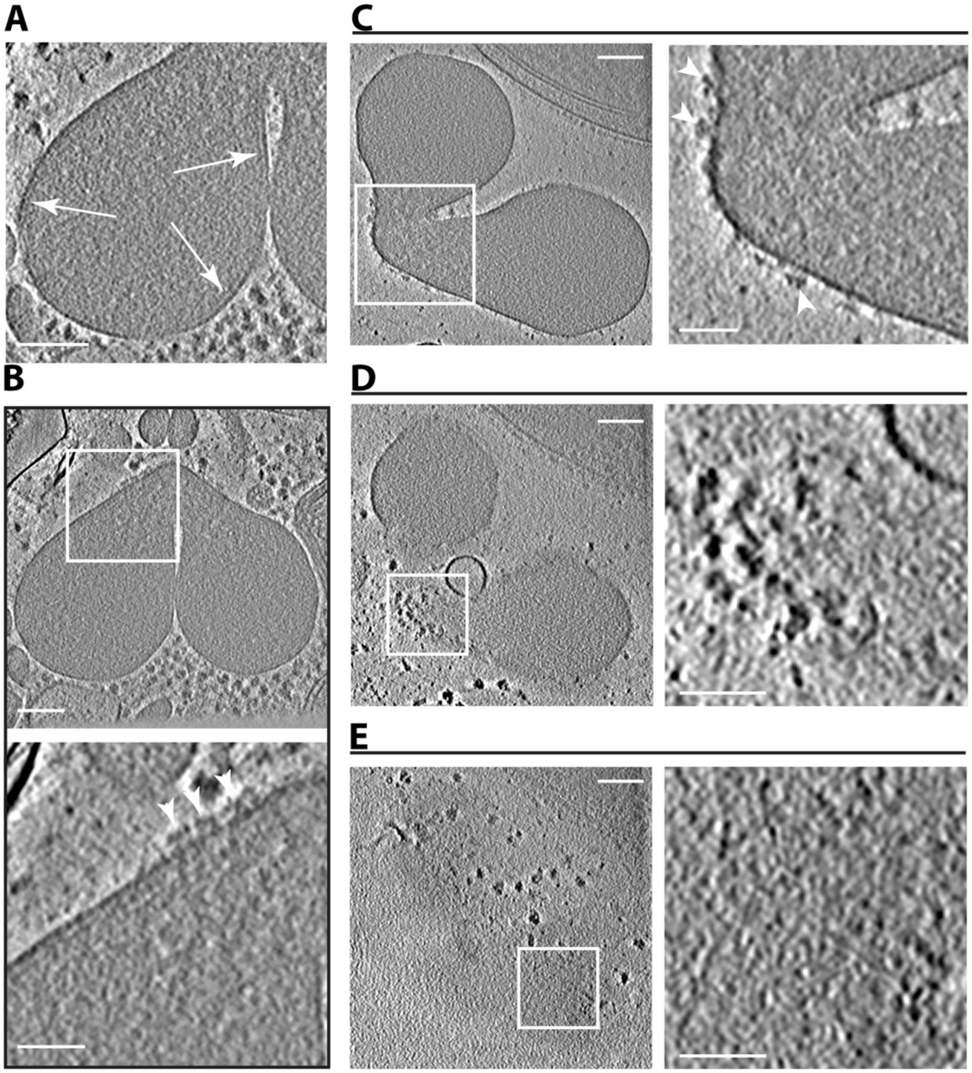


**Fig. S1. Distinct densities associated with rhoptry peripheries.**
(**A**) A tomographic slice showing a rhoptry with arrows pointing to a layer of continuous density reproducibly observed to line much of the organelle's limiting membrane. Scale bar, 100 nm. (**B**) Top panel is a tomographic slice showing the rhoptries. Scale bar, 100 nm. Bottom panel is zoomed in view of the square showing densities coating the cytosolic face of the membrane at the rhoptry's anterior end (arrowheads). Scale bar, 50 nm. (**C**) As for (B) except from a different tomogram. The arrowheads in the zoomed in view point to the densities coating the anterior end of the rhoptries. (**D**) A different slice of the same tomogram in (C) showing a surface view of the rhoptry neck region and the densities coating it (zoomed in view). (**E**) As for (D) except showing a surface view of the rhoptry bulbs. Note that the small associating densities apparent at the rhoptry necks are not apparent on the bulbs (zoomed in view). Scale bars, 100 nm or 50 nm for the zoomed in view.

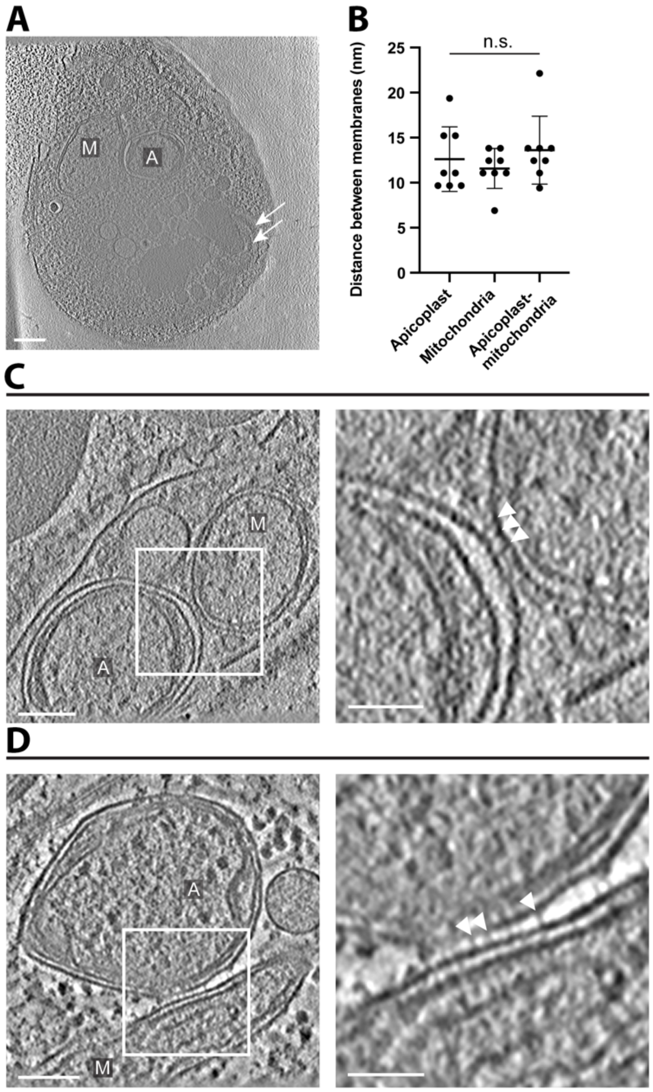


**Fig. S2. The apicoplast and mitochondrion maintain an intimate association in the merozoite cell.**

(**A**) A tomographic slice of a whole merozoite showing a mitochondrion (M) and apicoplast (A). The arrow marks the apical rings. Scale bar, 200 nm. (**B**) The distance between membranes was measured along a line profile plot of the inverted pixel values in the region where the two organelles are in closest apposition across. The plot shows the distance between the outermost two membranes of the apicoplast ("Apicoplast"), the inner and outer membranes of the mitochondrion ("Mitochondrion"), as well as between the limiting membranes of the two organelles ("Apicoplast-mitochondrion"). Apicoplast mean ± SD =12.6±3.6 nm, N=8; Mitochondrion mean ± SD =11.6±2.2 nm, N=8; Apicoplast to mitochondrion mean ± SD =13.6±3.8 nm, N=8; There was no significant difference between the groups, by one-way ANOVA. (**C**) Left panel- a tomographic slice showing a mitochondrion (M) and apicoplast (A). Right panel- a zoomed in view of the square in the left panel showing the area with minimum distance between the mitochondrion and the apicoplast with densities in the interface between them (arrowheads). Scale bars, 100 nm and 50 nm for the left and right panels, respectively. (**D**) As for (C), except from a different tomogram.

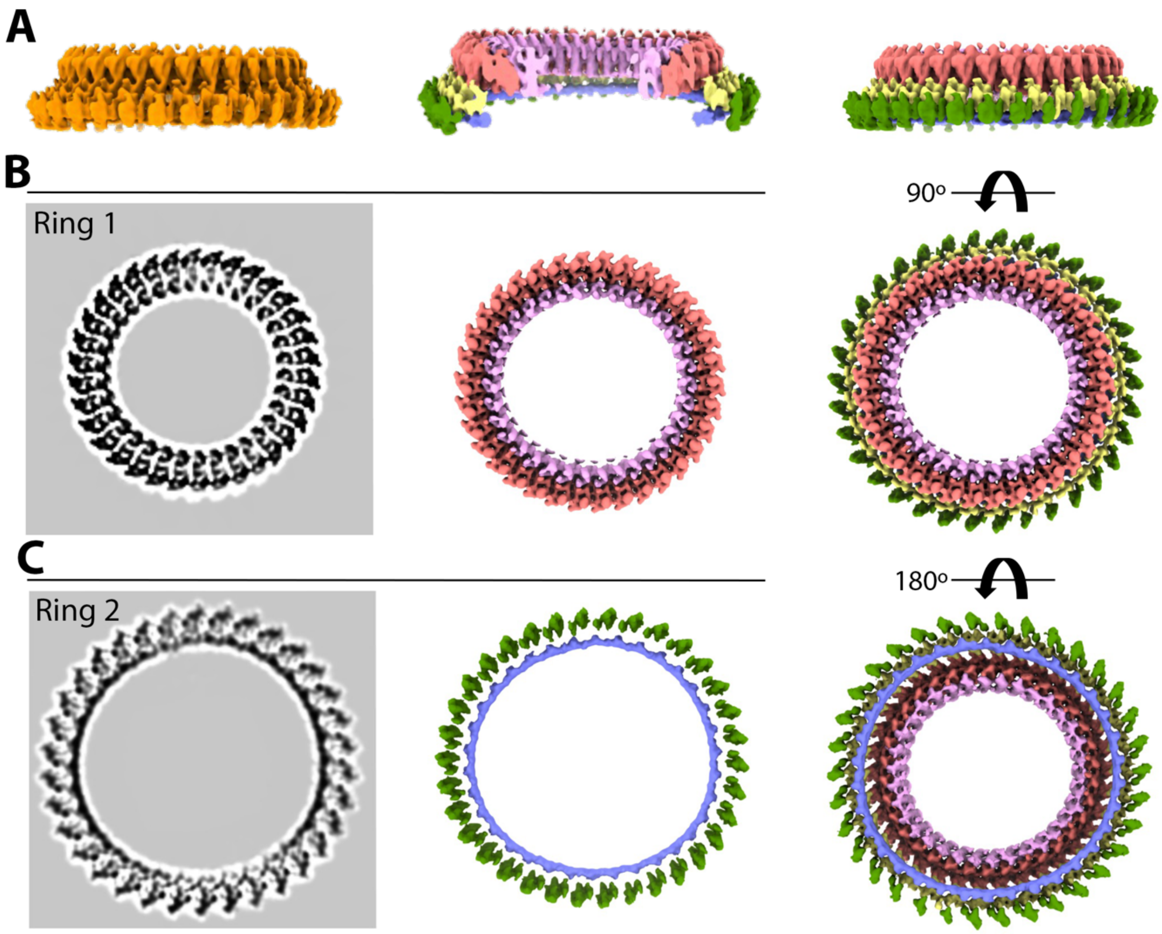


**Fig. S3. Rings 1 and 2 of *Plasmodium* show prominent 34-fold symmetry.**
(**A**) A side view of *Plasmodium* rings 1 and 2 of are shown in gold. The segmented cut view of the rings shows the inner and outer densities of Ring 1 in pink and red color, respectively and the inner and outer densities of ring 2 in blue and green color. The yellow density bridges the outer densities of ring 1 and ring 2. (**B, C**) The segmented cryo-EM map of Rings 1 and 2 for the tomogram in (A) are shown from a top (B) and bottom (C) view revealing 34 repeating units. Color scheme as in (A).

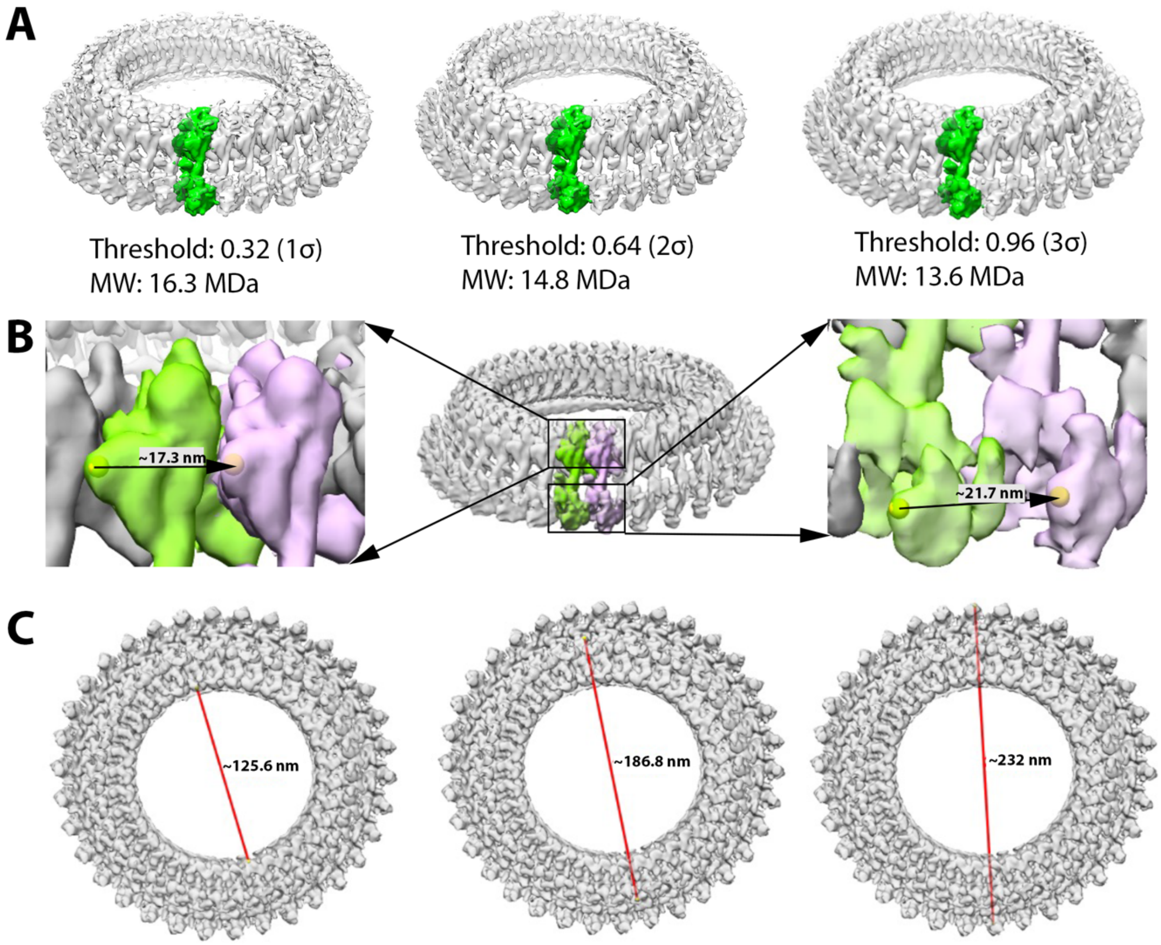


**Fig. S4. Measurements of *Plasmodium* rings 1 and 2.**
(**A**) A single repeating unit of ring 1 and 2 was segmented in green at three different thresholds. Its molecular size is estimated to be ~16.3MDa at thresholds 1σ, 14.8 MDa at 2σ, and 13.6 MDa at 3σ. (**B**) The spacing between neighboring units (green and pink) is ~17.3 nm on the outer edge of top ring 1 and ~21.7 nm on the outer edge of bottom ring 2. (**C**) The diameter of Rings 1 and 2 based on the averaged map. The inner and outer outlines of Ring 1 are ~125.6 nm and 186.8 nm, respectively. The outer diameter of Ring 2 is ~232 nm.

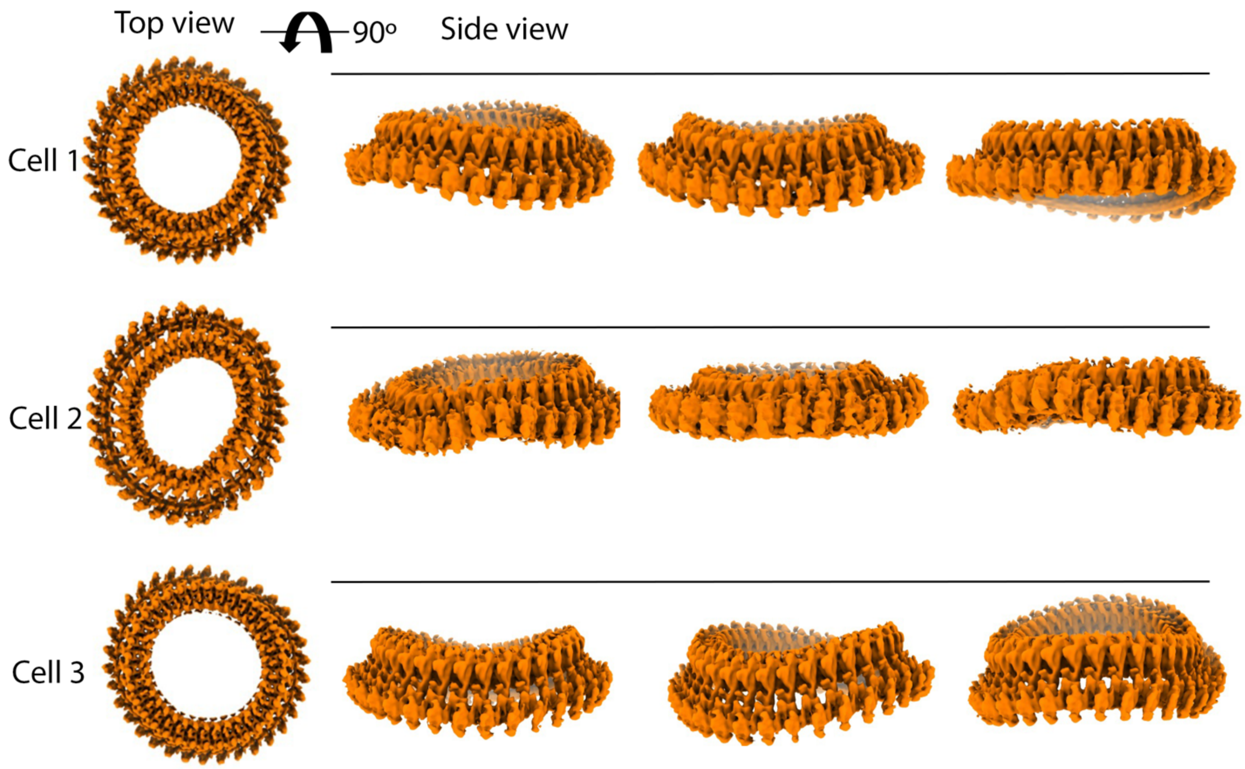


**Fig. S5. Views of *Plasmodium* rings 1 and 2 from different cells.**
The averaged map of rings 1 and 2 was mapped back to three tomograms (each representing a different cell). The different views of the rings showing twisting and bending in different ways suggests that the rings are flexible and the linkage between ring units is strong enough to withstand these torsional forces.

**Video 1. Tomogram showing a *Plasmodium falciparum* free merozoite with annotation of key subcellular structures.**

**Video 2. An apical vesicle is located underneath the rosette.**

**Video 3. The rosette can associate directly with the rhoptry neck region.**

**Video 4. The apical polar ring of Plasmodium presents a prominent 34-fold symmetry.**

**Video 5. The apical polar rings of Plasmodium merozoites and *Toxoplasma* tachyzoites show a similar organization and repeating pattern.**
